# Tunable Fibrin-Alginate Interpenetrating Network Hydrogels to Guide Cell Behavior

**DOI:** 10.1101/817114

**Authors:** Charlotte E. Vorwald, Tomas Gonzalez-Fernandez, Shreeya Joshee, Pawel Sikorski, J. Kent Leach

**Affiliations:** Department of Biomedical Engineering, University of California, Davis, Davis, CA, 95616; Department of Physics, Norwegian University of Science and Technology (NTNU), Trondheim, Norway; Department of Orthopaedic Surgery, UC Davis Health, Sacramento, CA 95817

**Keywords:** Interpenetrating network, endothelial cell, mesenchymal stromal cell, fibrin, alginate

## Abstract

Hydrogels are effective platforms for use as artificial extracellular matrices, cell carriers, and to present bioactive cues. Two common natural polymers, fibrin and alginate, are broadly used to form hydrogels and have numerous advantages over synthetic materials. Fibrin is a provisional matrix containing native adhesion motifs for cell engagement, yet the interplay between mechanical properties, degradation, and gelation rate is difficult to decouple. Conversely, alginate is highly tunable yet bioinert and requires modification to present necessary adhesion ligands. To address these challenges, we developed a fibrin-alginate interpenetrating network (IPN) hydrogel to combine the desirable adhesion and stimulatory characteristics of fibrin with the tunable mechanical properties of alginate. We tested its efficacy by examining capillary network formation with entrapped co-cultures of mesenchymal stromal cells (MSCs) and endothelial cells (ECs). We manipulated thrombin concentration and alginate crosslinking density independently to modulate the fibrin structure, mesh size, degradation, and biomechanical properties of these constructs. In IPNs of lower stiffness, we observed a significant increase in total cell area (1.72×10^5^ ± 7.9×10^4^ μm^2^) and circularity (0.56 ± 0.03) compared to cells encapsulated in stiffer IPNs (3.98×10^4^ ± 1.49×10^4^ μm^2^ and 0.77 ± 0.09, respectively). Fibrinogen content did not influence capillary network formation. However, higher fibrinogen content led to greater retention of these networks confirmed *via* increased spreading and presence of F-actin at 7 days. This is an elegant platform to decouple cell adhesion and hydrogel bulk stiffness that will be broadly useful for cell instruction and delivery.

## INTRODUCTION

Hydrogels are widely utilized in the pharmaceutical, biotechnology, and biomedical industry due to their high water content that mimics native tissues, biocompatibility, cell-friendly nature, and efficacy. Delivered as injectables, wound dressings, or implantable materials, hydrogels exhibit increased translational potential compared to other polymeric systems. Hydrogels derived from natural polymers are particularly promising, as they present native adhesion ligands to support cell interaction with the material. Fibrin is one broadly used natural polymer since it is abundant during the innate, wound healing process [1]. Fibrin exhibits both vasculogenic and anti-inflammatory potential for wound healing and tissue repair [2, 3]. Although this material enables cell infiltration and remodeling [4], it is relatively compliant and difficult to handle when delivered *via* implantation. Current formulations used in the clinic involve fibrin clotting components at supraphysiological doses, resulting in fast gelation times that add deleterious stress on encapsulated cells.

Hydrogels are composed of polymer chains frequently comprised of one repeating monomer, recognized as homopolymers [5]. While such polymer chain formulations can be characterized relatively easily, their bulk properties are limited to these homotypic interactions. Copolymers facilitate diversity in hydrogel formulations, yet these materials are constrained by similar disadvantages, as polymer composition remains a limiting factor for hydrogel properties [6]. Interpenetrating networks (IPNs) are created with two or more distinct polymer strands interlaced among each other [7] and represent an exciting platform to address these shortcomings. Each polymer strand is crosslinked through distinct mechanisms, either simultaneously or in parallel. These formulations more broadly capture the potential of each polymer network and maximize tunability [8]. Among polymers used in IPNs, alginate is commonly studied due to its tunable, robust mechanical properties. IPNs have been formed by combining collagen-alginate [9, 10], basement membrane-alginate [11], fibrin-alginate [12], and others. Previous studies have confirmed the potential to combine two polymers and modulate both adhesion and mechanical characteristics in hydrogel constructs. Yet, there is limited knowledge regarding the interplay between crosslinking mechanism, subsequent changes in biophysical properties of IPNs, and resulting cell response.

Herein, we describe the fabrication and characterization of a fibrin-alginate IPN to balance the compliance of fibrin with the mechanical stability and tunability of alginate. We report a unique method to create these constructs that retains fibrin structure within a dense alginate network. We utilized light reflection microscopy, a non-invasive, label-free imaging modality, to confirm changes in fibrin structure and evaluate IPN structural properties. We used clinically relevant cell sources to test the efficacy of these IPNs, namely endothelial cells (ECs) and mesenchymal stromal cells (MSCs), which synergistically promote wound healing through capillary network formation [13]. EC-MSC response was dependent on alginate crosslinking density and fibrinogen content. This work provides insight into the interplay of mechanical properties and cell spreading on capillary formation and retention and establishes FA IPN as a promising biomaterial for biomedical applications.

## MATERIALS AND METHODS

### IPN synthesis

UltraPure MVG sodium alginate (Pronova, Lysaker, Norway) was oxidized to a degree of oxidation of 1% with sodium periodate (MilliporeSigma, St. Louis, MO) as previously described [14] and dissolved in PBS at 3% w/v. Fibrinogen (MilliporeSigma) was mixed with alginate solution at 30 mg/mL and allowed to dissolve at 37°C in PBS overnight. This solution remained sterile and stored at 4°C until use for subsequent experiments.

3.5 kDa MWCO dialysis membranes (Spectrum Labs, Rancho Dominguez, CA) were hydrated in sterile ultrapure water for at least 10 min. We prepared solutions of calcium chloride (CaCl_2_, MilliporeSigma) in water at 10 mM or 40 mM. To obtain a resulting 2% w/v alginate and 20 mg/mL fibrinogen IPN, 66 μL of fibrinogen-alginate solution was mixed with 33 μL of thrombin solution with a concentration range of 0-75 U/mL, and 80 μL was immediately pipetted into 8 mm circular, silicon molds. Fibrin networks were allowed to form for 45 min at 37°C. A hydrated dialysis membrane was then placed gently on top of molds, and CaCl_2_ solution was pipetted on top until fully covered. Alginate was allowed to crosslink for 30 min at 37°C. Resulting constructs were gently transferred from molds to non-coated 8-well chamber slides (ibidi, GmbH, Planegg, Germany).

### Fibrin characterization *via* light reflection and confocal microscopy

66 μL of Fibrinogen-Alginate solution was mixed with 33 μL of thrombin (75 U/mL, 7.5 U/mL, and 0.75 U/mL) and pipetted directly into non-coated 8-chamber wells (ibidi) for a resulting fibrinogen and thrombin concentration of 20 mg/mL and 25 U/mL, 2.5 U/mL, and 0.25 U/mL, respectively. Fibrin was allowed to form for 45 min at 37°C. Reflection confocal microscopy was performed using the Leica TCS SP8 (Leica, Wetzlar, Germany) at 40X magnification to observe fibrin structure within FA IPN constructs.

### Gelation kinetics measurements of fibrin in alginate

66 μL of Fibrinogen-Alginate solution was mixed with 33 μL of thrombin (75 U/mL, 7.5 U/mL, and 0.75 U/mL), and 50 μL was pipetted directly into each well of a flat, clear 96-well plate. Optical density was measured at 350 nm and 550 nm in triplicate at 4 min intervals for 45 min at 37°C. Similar trends from 350 nm and 550 nm readings ensured that results were due to light scattering and not due to absorption [15, 16]. Fibrin with respective thrombin concentrations served as positive controls. PBS, IPNs without thrombin, and fibrinogen without thrombin served as negative controls.

### Characterization of IPN physical properties

Viscoelastic properties of acellular IPNs were measured on a Discovery HR-2 hybrid stress-controlled rheometer (Thermal Analysis Instruments, New Castle, DE) equipped with an 8 mm parallel plate geometry. Gels were tested at a strain of 0.5% and frequency sweep from 0.1 to 10 rad/s. The storage modulus was reported at 1.58 rad/s.

Mesh size, *r*_*mesh*_, was calculated from data based on rheological measurements [17] and approximated by the following equation:

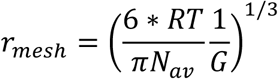

where R is the gas constant, T is the absolute temperature, N_av_ is Avogadro’s number, and G is the storage modulus. Initial storage modulus of crosslinked gels was measured, and gels were subsequently weighed to obtain wet weight. Constructs were then frozen at −80°C overnight, lyophilized for 1 day, and weighed again to obtain dry weight.

### Cell culture

Human cord blood-derived endothelial colony forming cells were kindly provided by Prof. Eduardo Silva (UC Davis) and expanded in EGM-2 supplemented media (PromoCell, Heidelberg, Germany) with gentamycin (50 µg/mL; Invitrogen) and amphotericin B (50 ng/mL; Invitrogen) under standard conditions (37°C, 5% CO_2_, 21% O_2_) until use at passage 5 [18]. Human bone marrow-derived MSCs (Lonza, Walkersville, MD) from a single donor (22-year-old male) were expanded without further characterization in growth medium (GM) consisting of minimum essential alpha medium (α-MEM; Invitrogen, Carlsbad, CA) supplemented with 10% fetal bovine serum (FBS; Atlanta Biologicals, Flowery Branch, GA) and 1% penicillin/streptomycin (Gemini Bio-Products, Sacramento, CA). MSCs were cultured under standard conditions until use at passage 5. Media changes were performed every 2-3 days. For each experiment, aliquots were derived from the same batch of serum to ensure consistency.

### EC-MSC encapsulation in FA IPN constructs

ECs and MSCs were suspended in Fibrinogen-Alginate solution at 1×10^6^ cells/mL and 2×10^6^ cells/mL, respectively, for a total concentration of 3×10^6^ cells/mL. 66 μL of Fibrinogen-Alginate cell suspension was mixed with 33 μL of thrombin (7.5 U/mL), and 80 μL was pipetted directly into 8 mm silicon molds. Fibrin was allowed to form for 45 min at 37°C. Meanwhile, a hydrated dialysis membrane was placed gently on top of molds, and CaCl_2_ solution was added on top until fully covered. Alginate was allowed to crosslink for 30 min at 37°C. Resulting constructs were gently transferred from molds to non-coated 8-chamber slides (ibidi) and cultured in 250 μL 3:1 EGM-2: α-MEM under standard conditions. Media was refreshed every 48 hours.

### Evaluation of cell spreading within IPNs using confocal microscopy

Before encapsulation, ECs were stained with CellTrace™ Oregon Green® 488 (carboxy-DFFDA SE) (Invitrogen) and MSCs were stained with CellTrace™ Far Red Cell Proliferation Kit (Invitrogen). Confocal microscopy was performed using the Leica TCS SP8 (Leica) on day 1, 3, and 7 of culture. Images were processed and analyzed in ImageJ (NIH, Bethesda, MD). Confocal images were converted to 500 × 500 pixel size, converted to binary, and circularity and area of cells, with an area of at least 50 μm^2^, well below the average size of ECFCs [19], were measured. Values are reported by cell population channel acquired for each material group. At day 7 of culture, gels were fixed with 4% paraformaldehyde at 4°C overnight, washed twice with PBS, and permeabilized with 0.05% Triton-X 100 for 5 min at room temperature. Gels were stained with Alexa Fluor 488 Phalloidin solution (Thermo Fisher; 1:40 in PBS) and incubated at room temperature for 1 hr. Gels were washed twice with PBS and subsequently imaged using confocal microscopy.

### Statistical analysis

Data are presented as means ± standard deviation. Statistical significance was assessed by either one-way ANOVA, two-way ANOVA with Tukey’s multiple comparisons test, or Student’s t-test when appropriate. *p*-values <0.05 were considered statistically significant. Statistical analysis was performed using GraphPad Prism® 8 analysis software (GraphPad Software, La Jolla, CA). Different letters denote statistical significance between groups, while data sharing a letter are not statistically different from one another.

## RESULTS

### Fibrin structure and gelation of IPNs are dependent on thrombin concentration

The formation of uniform, well-formed fibrin fibers within the alginate solution before alginate crosslinking occurs was a key design criterion for successful IPN formulation, as this is representative of mature, fibrin fibers found in blood clots [20]. To determine the interplay of thrombin concentration and fibrin fiber morphology, we assessed fibrin structure within crosslinked alginate using confocal reflection microscopy. We observed a striking increase in fibrin thickness and reduced alignment as thrombin concentration increased (**Fig. 3A**). Fibrin control gels exhibited decreased mesh size with increasing thrombin concentrations. Fibrin structure for gels crosslinked with both 0.25 and 2.5 U/mL thrombin possessed a consistent mesh size throughout the gel. However, fibrin gels made with 25 U/mL thrombin possessed a dense, fine mesh with large, open pockets. We observed more open mesh structure with thicker fibers in IPNs compared to fibrin controls of corresponding thrombin concentration. In IPNs, we also observed increased density of fibrin structures as thrombin concentration increased.

**Figure 1.**
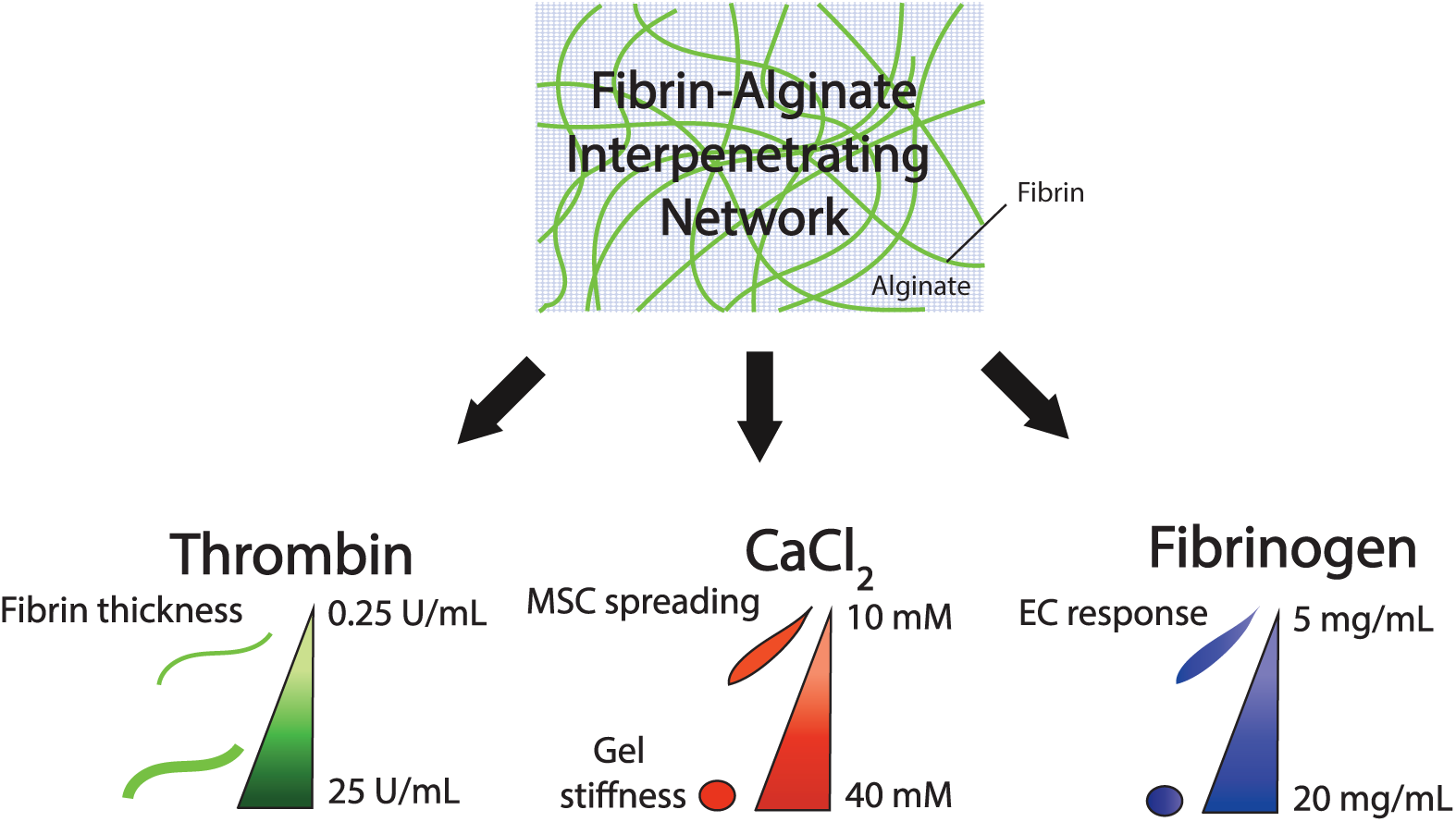
Schematic of parameters used to tune biophysical properties and characteristics of FA IPNs. Tuning of crosslinking components affects polymer structure, mesh density, biomechanical properties, and cell response.

**Figure 2.**
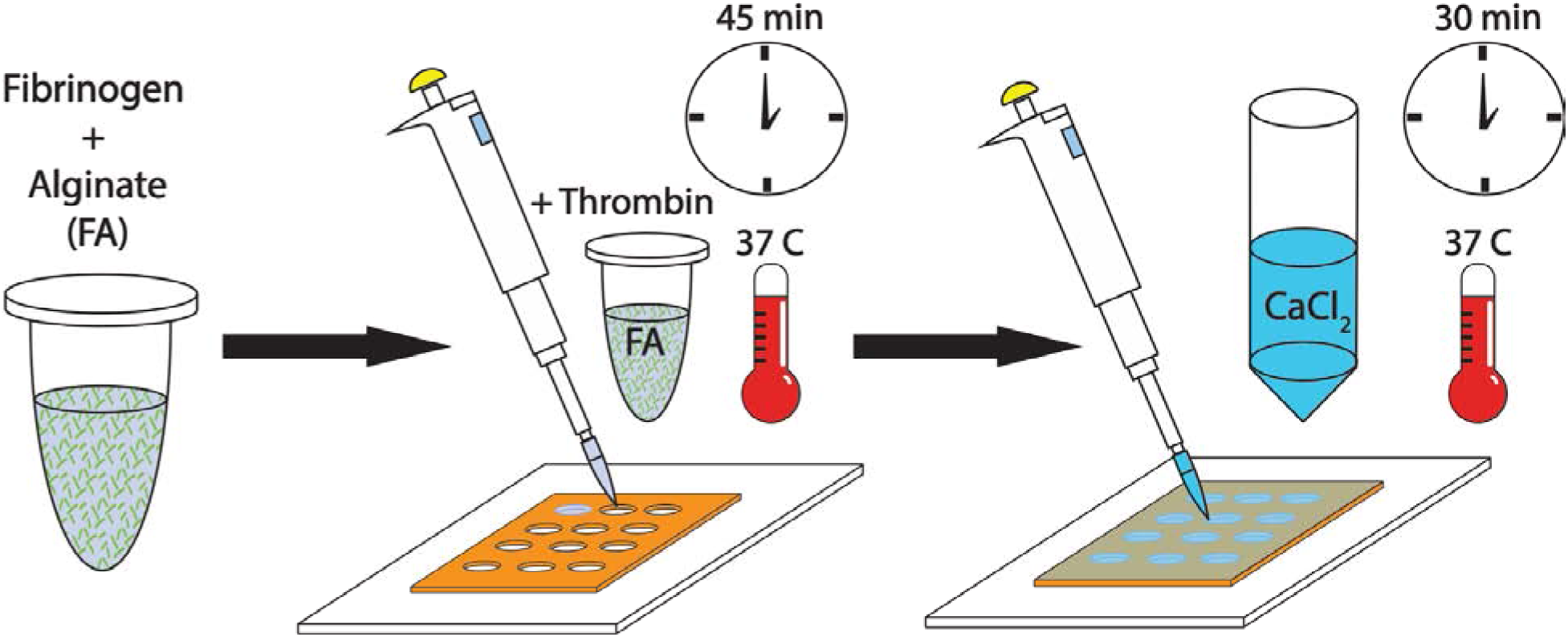
Experimental outline of FA IPN fabrication. Fibrinogen and alginate are combined to generate FA IPNs. Independent crosslinking mechanisms are promoted through the addition of fibrin and CaCl_2_.

**Figure 3.**
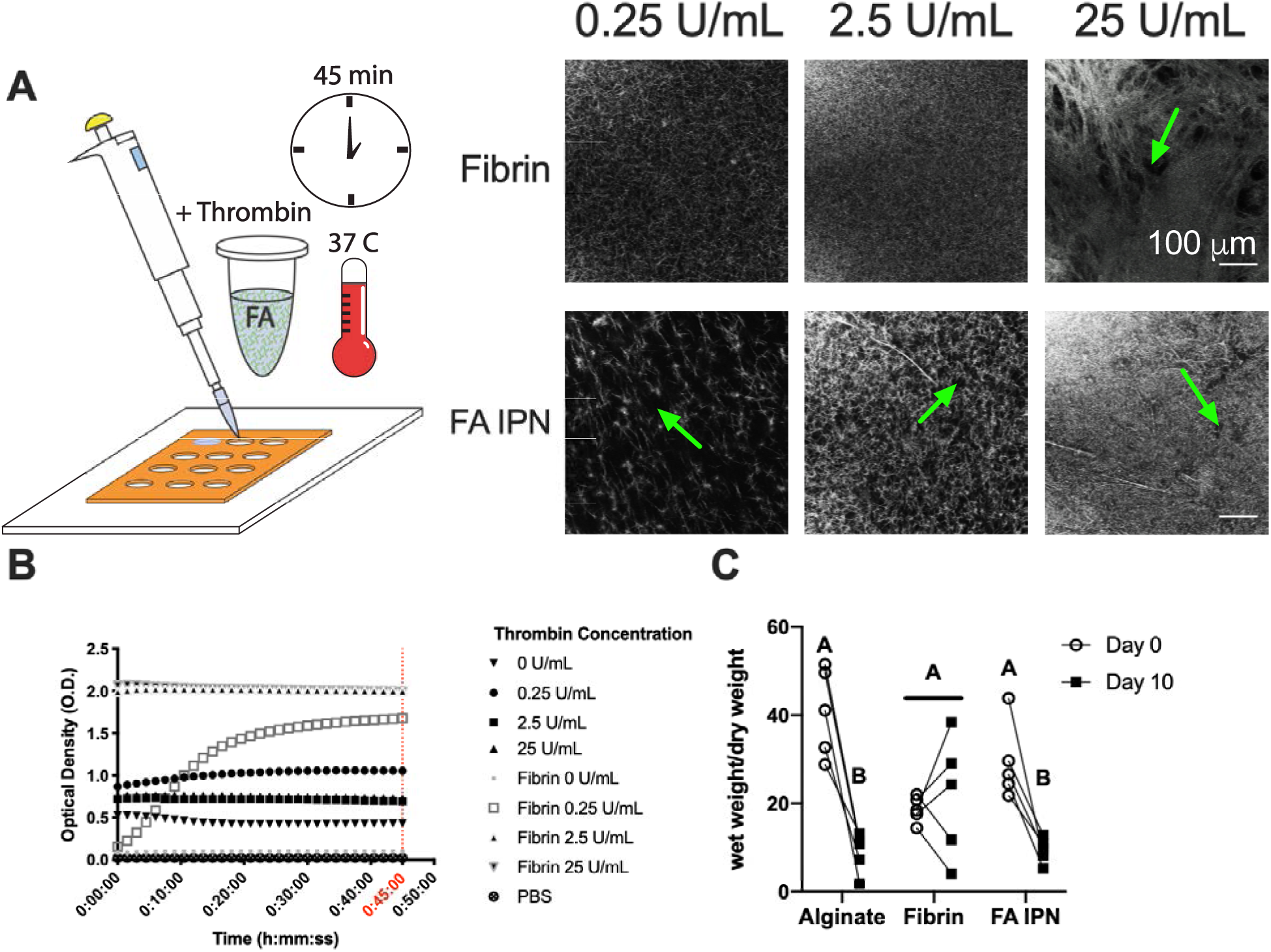
Fibrin structure and gelling kinetics in alginate are dependent upon thrombin concentration. **(A)** Schematic and representative images of fibrin structure investigated using light reflection microscopy with varying thrombin concentrations. Green arrows indicate the presence of open pockets within the IPN. **(B)** Optical density of fibrin constructs at 550 nm (n=4-5). **(C)** Wet weight normalized to dry weight of constructs cultured over 10 days (n=5). Different letters denote statistical significance (*p*<0.05).

We measured gel turbidity during the gelation process to explore the kinetics of gelation in IPNs (**Fig. 3B**). Turbidity, and as a consequence optical density (O.D.), increased as the fibrin fibers were formed [15, 16]. We compared fibrin formation in fibrin only controls and fibrin-alginate solutions. At 45 minutes, PBS and fibrin without thrombin exhibited minimal O.D. (0.03 ± 0.001 and 0.1 ± 0.003, respectively), as expected. IPNs without thrombin exhibited higher initial and resulting O.D. (0.43 ± 0.29) compared to both PBS and fibrin without thrombin. O.D. increased for fibrin gels as thrombin concentration increased. Fibrin with 0.25 U/mL thrombin showed a slow increase in O.D. over time, while fibrin gels made with 2.5 and 25 U/mL thrombin demonstrated a striking increase in overall O.D. values over 45 minutes. Similarly, O.D. increased in IPNs with increasing thrombin concentrations. However, IPN O.D. was highest with 0.25 U/mL thrombin (1.06 ± 0.14) compared to 2.5 U/mL (0.69 ± 0.29) and 25 U/mL thrombin (0.73 ± 0.52).

To understand the contribution of fibrin structure on degradation kinetics of FA IPNs, fibrin and alginate were fully crosslinked with 2.5 U/mL thrombin and 10 mM CaCl_2_, respectively, and we measured wet and dry weight over 10 days (**Fig. 3C**). We observed a significant decrease in normalized wet weight between day 0 and day 10 for alginate (40.7 ± 10.0 to 8.9 ± 4.5, *p*=0.0002) and FA IPN groups (29.2 ± 8.67 to 9.4 ±2.9, *p*=0.0012), but we observed similar weights in fibrin controls between timepoints (17.6 ± 18.6 and 21.5 ± 13.7, *p*=0.66). On day 0, alginate possessed a greater normalized wet weight (40.8 ± 10.0) compared to fibrin control (18.6 ± 2.9) but reached similar values by day 10. IPNs exhibited similar trends on day 0 and day 10 compared to alginate controls.

### Mesh size and mechanical properties are tunable *via* calcium chloride crosslinking

We characterized overall construct properties after alginate crosslinking (**Fig. 4A-B**). To demonstrate that IPN constructs can be molded into desired morphologies, we formed gels as previously described but with molds cut with distinct edges. Distinct shapes cannot be discerned in fibrin gels that lose their form upon handling and transfer (**Fig. 4B**). Rheological measurements (**Fig. 4C**) revealed similar storage moduli between alginate and FA IPNs crosslinked with 10 mM CaCl_2_ (627.8 ± 100.5 and 831.0 ± 205.1 Pa) compared to fibrin controls (250.4 ± 74.0 Pa). Alginate and FA IPNs crosslinked with 40 mM CaCl_2_ possessed significantly increased storage moduli (3144.3 ± 686.4 and 2587.6 ± 1266.3 Pa, *p*=0.0001) compared to alginate and FA IPNs crosslinked with 10 mM CaCl_2_ and fibrin controls. Average mesh size was inversely correlated with storage modulus (**Fig. 4D**). Mesh size decreased in gels when crosslinked with 10 mM CaCl_2_ versus 40 mM CaCl_2_ in alginate (3.5 ± 0.2 nm vs. 2.0 ± 0.1 nm, *p*=0.002). FA IPNs exhibited a similar trend, with decreased mesh size as CaCl_2_ concentration increased (3.2 ± 0.3 nm and 2.3 ± 0.4 nm, *p*=0.0027). Mesh size in IPNs or alginate gels were smaller than fibrin gel controls (4.8 ± 0.6 nm; *p*≤0.0001).

**Figure 4.**
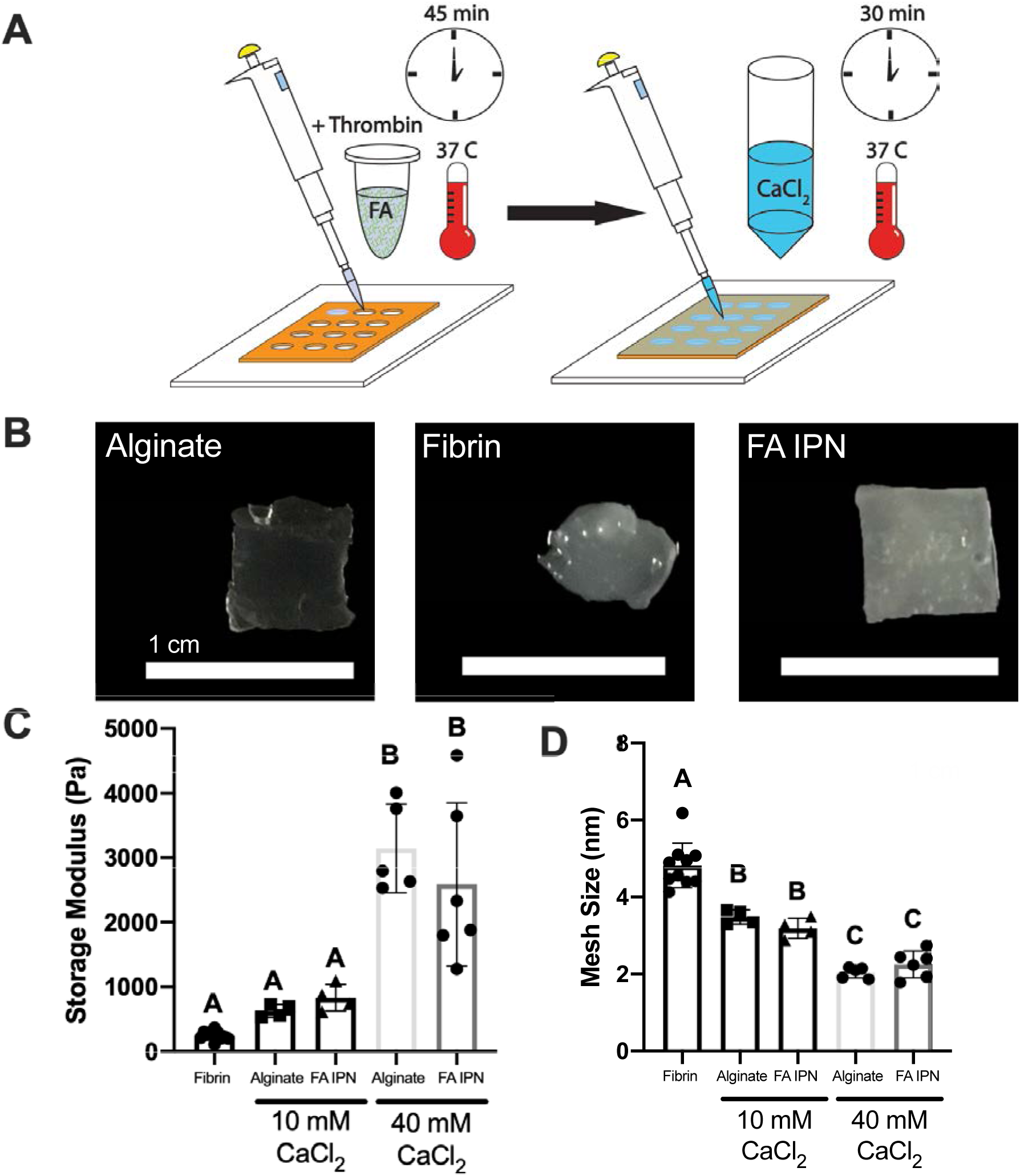
Material characterization of IPNs as a function of calcium chloride concentration. **(A)** Schematic of FA IPN fabrication. (**B**) Images of resulting constructs after 30 min of gelation with 40 mM CaCl_2_ and transfer. (**C**) Storage modulus and **(D)** calculated mesh size of FA IPNs with varying calcium chloride concentration (n=4-10). Different letters denote statistical significance (*p*<0.05).

### EC-MSC spreading is a function of alginate crosslinking density

We aimed to understand the contribution of fibrin and alginate crosslinking on EC and MSC spreading within IPNs. On day 3 of co-culture, EC and MSC spreading was a function of CaCl_2_ concentration, with greater spreading observed in IPNs formed with 10 mM CaCl_2_ compared to 40 mM CaCl_2_ (**Fig. 5A**). Cells in alginate controls, which lacked adhesion motifs, exhibited spherical morphology regardless of CaCl_2_ content, while fibrin controls supported cell spreading as expected. Quantification of total area and circularity of cells on day 3 of co-culture (**Fig. 5B**) support these qualitative observations. Quantification of total area revealed greater cell area in fibrin (1.39×10^5^ ± 4.1×10^4^ μm^2^) compared to alginate crosslinked with 10 mM CaCl_2_ (5.50×10^4^ ± 3.4×10^4^ μm^2^; *p*=0.02). FA IPNs crosslinked with 10 mM CaCl_2_ exhibited similar total cell area (1.72×10^5^ ± 7.9×10^4^ μm^2^) compared to fibrin, yet cell area in IPNs crosslinked with 40 mM CaCl_2_ was significantly reduced (3.98×10^4^ ± 1.49×10^4^ μm^2^; *p*=0.001). Cell areas within alginate gels crosslinked with either 10 mM CaCl_2_ (5.51×10^4^ ± 3.42×10^4^ μm^2^) or 40 mM CaCl_2_ (7.44×10^4^ ± 1.51×10^4^ μm^2^) were comparable. Circularity revealed overall cell shape of cell populations within these gels, with circularity of 1.0 representing cells with symmetrical circular properties. Cells in fibrin controls (0.46 ± 0.02) had lower circularity versus alginate controls crosslinked with 10 mM CaCl_2_ (0.79 ± 0.11; *p*=0.0001) and 40 mM CaCl_2_ (0.82 ± 0.04; *p*=0.0001). IPNs crosslinked with 10 mM CaCl_2_ (0.56 ± 0.03) resulted in intermediate circularity values between fibrin and alginate controls, while IPNs crosslinked with 40 mM CaCl_2_ (0.77 ± 0.09) induced similar circularity values to both alginate controls. To understand whether fibrin networks within these constructs may be instructing these differences, we observed fibrin structure on day 3 of co-culture with light reflection microscopy (**Fig. 5C**). As expected, we did not observe fiber structures within alginate controls with light reflection, while dense, mesh structures were observed throughout fibrin gels encapsulating both cell types. In IPN hydrogels, we observed similar fibers, although less dense compared to fibrin controls. Furthermore, fibrin fibers were more apparent in IPNs crosslinked with 40 mM CaCl_2_ compared to those with 10 mM CaCl_2_.

**Figure 5.**
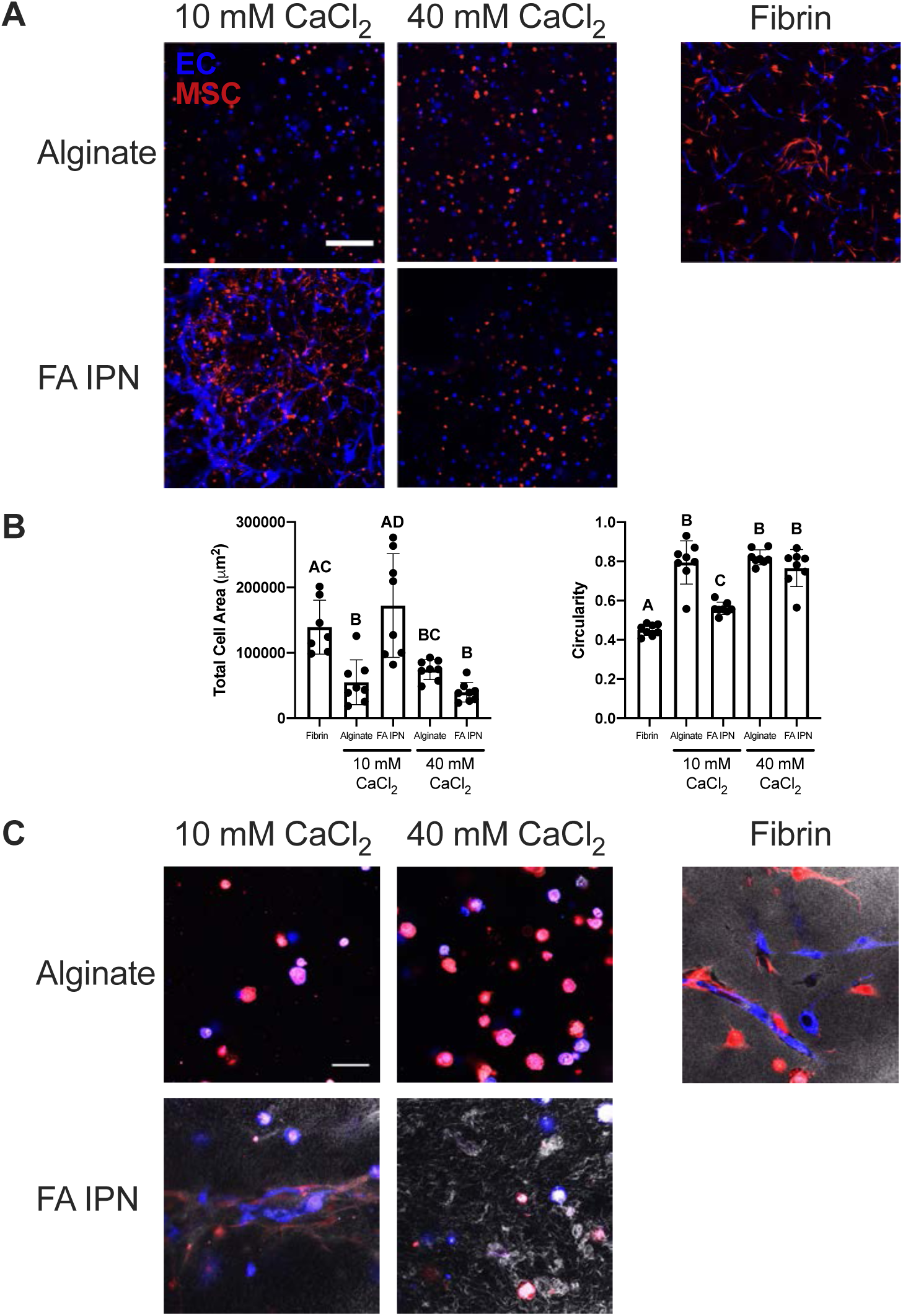
Alginate crosslinking density affects EC-MSC response *in vitro.* **(A)** Confocal images of EC (violet) and MSC (red) on day 3 of culture. Scale bars are 250 μm. **(B)** Respective quantification of total cell area and circularity at day 3 (n=7-8). **(C)** Confocal images of fibrin structure (grayscale) with EC (violet) and MSC (red) at day 3. Scale bars are 50 μm. Different letters denote statistical significance (*p*<0.05).

### EC network retention is a function on fibrinogen content within FA IPNs

Since alginate crosslinking density affects the degree of EC and MSC spreading, we aimed to investigate the role of fibrin on network formation when keeping crosslinking density constant. We formed IPNs with 10 mM CaCl_2_ while varying the amount of fibrinogen incorporated in the FA IPN: 5 mg/mL, 10 mg/mL, and 20 mg/mL. On day 3, we observed that EC network formation was inversely correlated with fibrinogen concentration in fibrin gels (**Fig. 6A-B**). We observed increases in cell area in fibrin controls compared to alginate controls and IPNs crosslinked with 40 mM CaCl_2_. Cell area was similar in FA IPNs crosslinked with 10 mM CaCl_2_ compared to fibrin controls, regardless of fibrinogen content. Measurement of cell circularity revealed a similar trend, with no dependence of circularity on fibrinogen concentration in fibrin gels or FA IPNs. However, network formation was maintained in groups with increased fibrinogen on day 7 (**Fig. 6C**). This retention of network formation is further supported by phalloidin staining to detect F-actin, which signifies stress fiber formation. FA IPNs with 5 mg/mL fibrinogen exhibited little F-actin expression. Cells entrapped in FA IPNs with 10 and 20 mg/mL fibrinogen exhibited increased F-actin expression, with increased spreading and distinct, stretched morphology of stress fibers.

**Figure 6.**
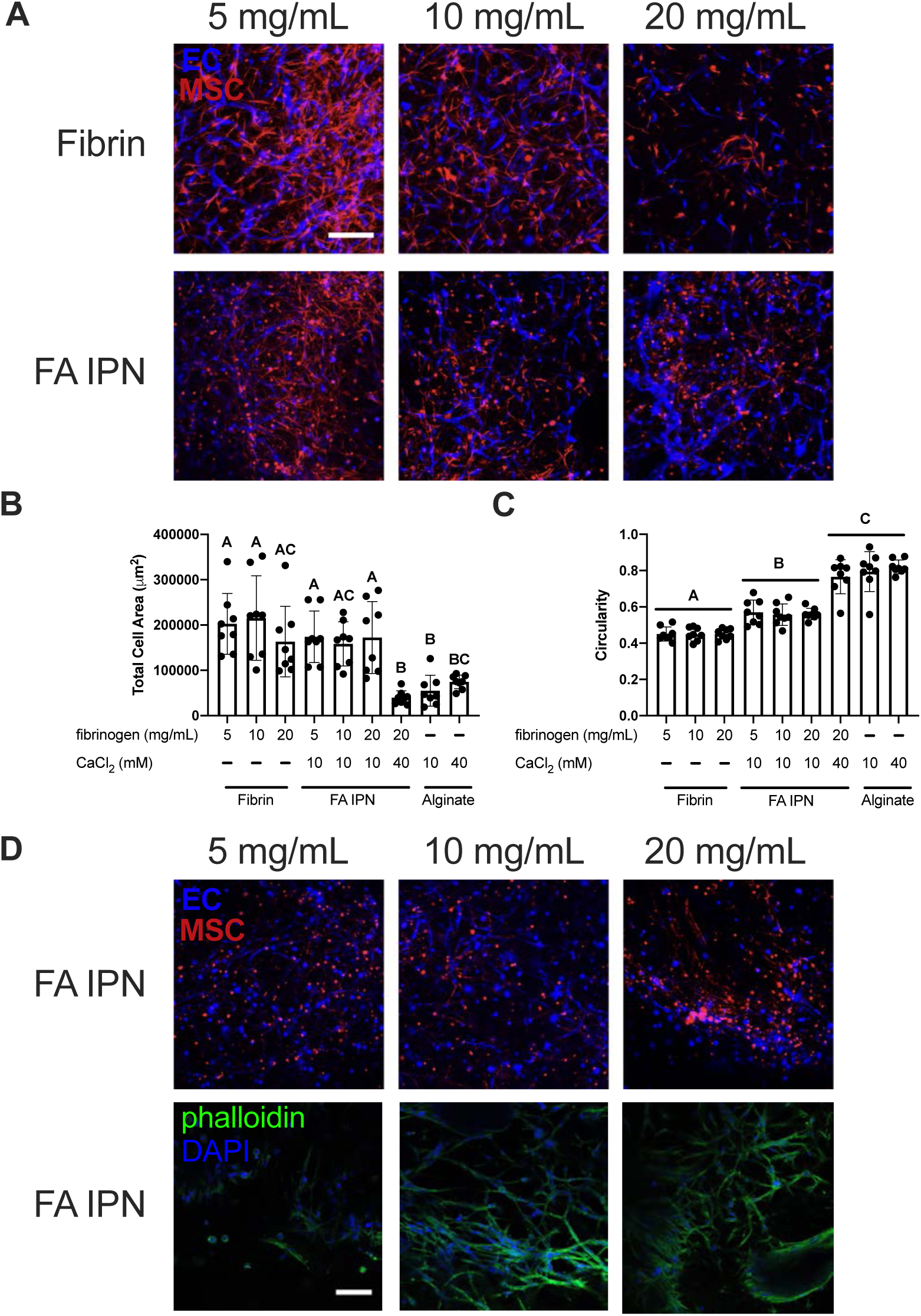
Fibrin content affects EC-MSC network formation and retention. **(A)** Confocal images of ECs (violet) and MSCs (red) in FA IPNs with varying fibrin content on day 3 of culture. Scale bars are 250 μm. **(B)** Quantification of total area and **(C)** circularity (1.0 indicates a perfect circle) of ECs and MSCs in field of view on day 3 of culture (n=8). **(D)** Confocal images of Ecs (violet) and MSCs (red) in FA IPNs (top) with respective phalloidin (green) and DAPI (violet) staining (bottom) on day 7. Scale bars are 250 μm. Different letters denote statistical significance (p<0.05).

## DISCUSSION

Hydrogels have tunable characteristics for adhesion, mechanical properties, and degradation, making them valuable platforms to mimic the native extracellular matrix (ECM). Standard hydrogel formulations are often formed of a single polymer, limiting their tunability as a function of composition. IPN hydrogels offer a strategy to achieve a preferred embodiment of biomaterial by decoupling function from composition [21]. Thus, IPNs have been formed from a number of materials including collagen-alginate [10], fibrin-alginate [12], and fibrin-hyaluronic acid [22]. However, formulations utilizing matricellular proteins such as collagen rely on the concentration of the polymer to drive mechanical and structural properties, while formulations using fibrin require one-step protocols for rapid crosslinking and gelation. In this work, we formulated and characterized a fibrin-alginate IPN to independently tune adhesivity and hydrogel physical properties, which was validated by testing its capacity to support network formation by an entrapped co-culture of MSCs and endothelial cells.

Alginate is one of the most broadly used hydrogels for biological applications because it possesses tunable mechanical properties, induces little inflammation, and is amenable to a variety of crosslinking methods [23]. As alginate is a bioinert polymer, it requires covalent modification with adhesive motifs to enable cell adhesion. This has been achieved with a variety of ECM components, with adhesive peptides such as Arginine-Glycine-Aspartic Acid (RGD) being the most common [24]. Peptides that are covalently bound to the polymer backbone promote adhesion but fail to mimic the complex nature of the ECM, while chemical modification of polymers with full sequence proteins is inefficient and may have interactions with other components that impair crosslinking. IPNs represent an exciting approach to address this limitation. For example, collagen-alginate IPNs have been investigated for tuning stromal cell morphology and trophic factor secretion. However, collagen content and structure are limited by protein concentration [25]. Fibrin has also been employed in IPNs, as various protein and crosslinking parameters can be adjusted for desired gelation strategies [26]. Fibrin-alginate IPNs were previously used as a niche for follicle development [12], in which fibrin structure was tailored to control trophic factor secretion. Fibrin provides natural binding motifs to allow adhesion and remodeling and has been extensively used in wound healing models [27], making it an ideal component for incorporation in this IPN.

During the native wound healing process, thrombin cleaves fibrinogen, resulting in a dense, polymerized fibrin network. We aimed to create a fibrin network within alginate that displays such thick, uniform fibers [20]. Thrombin concentration affects fiber thickness and density within fibrin gels [28, 29], with higher concentrations resulting in thinner, yet more dense fibers [30]. Within IPN hydrogels, we also observed distinct patterns of fibrin structure as a function of thrombin concentration. Fibrin structures in IPNs appeared thicker yet less compact compared to fibrin controls, most notably with 2.5 U/mL thrombin. We speculate that diffusion of thrombin is slower in the precursor fibrinogen-alginate solution compared to PBS due to increased viscosity. This is supported by our data on gelation kinetics, as IPNs exhibit reduced O.D. compared to respective fibrin controls. In all groups, we observed saturation in fibrin crosslinking through plateaus in gelation kinetics, suggesting that although thrombin dynamics may be different, the protein activity remains effective. These findings establish that manipulation of thrombin concentration is a novel method to influence fibrin arrangement within fibrin-alginate IPNs.

Although the availability of binding sites for cells is critical for cell attachment and function, the bulk properties of hydrogels play an equally important role. Alginate is widely used as a platform to study the effect of mechanical cues because the gelation [31, 32], swelling ratio [33], crosslinking density, and resultant moduli [34] can be easily tuned by calcium ion concentration. Thus, the degree of alginate crosslinking is another critical component for tuning cell response within this IPN. We observed increases in hydrogel storage modulus as the concentration of CaCl_2_ increased. Furthermore, alginate crosslinking density affected MSC and EC spreading, with the greatest cell spreading within IPNs crosslinked with 10 mM CaCl_2_. Compared to fibrin gels, we observed rapid degradation of IPNs and alginate controls, suggesting alginate is degrading in our system. We and others have emphasized the importance of construct degradation and proteolytic cleavage kinetics suitable for endothelial network formation [35]. The differences in IPN hydrogel bulk properties crosslinked with 10 mM or 40 mM CaCl_2_ were driven by multiple variables including degradation and mesh size. These two parameters work in concert to present adhesion sites, allow for solutes to diffuse through the polymer network, and ultimately create a dynamic environment. While these data stress the importance of mesh structure and alginate degradation on EC and stromal response in these polymer networks, it also reveals that alginate stiffness is a limiting factor of FA IPN integrity. Thus, other methods to increase stiffness are necessary such as increasing the amount of alginate in solution [6] or changing alginate properties [14] once cell networks are already formed.

In order to achieve our goal of developing a functional IPN, we established a novel method for adding fibrinogen to the polymer formulation, resulting in a means to tailor the adhesivity and bulk mechanical properties of a clinically relevant hydrogel. While we interrogated elements of mesh size to understand the role of structure on cell response, the applied analytical formula is limited in its complexity. The mesh structure between alginate and fibrin are significantly different, with alginate exhibiting mesh size in nanometer scale and fibrin in micrometer scale. Full investigation *via* imaging modalities such as atomic force microscopy could be used to reveal true structure morphology of alginate and fibrin in its hydrated form. Although simplified, our reports on mesh size highlight the interplay between structural and mechanical properties on tailoring cell response. Furthermore, outputs for cell response (area and circularity) are 2D in nature, not fully describing its 3D network capacity. Nonetheless, we believe clear conclusions can be drawn through these parameters to characterize how cells interact with the IPNs.

This fibrin-alginate IPN formulation is an elegant method to modulate cell adhesion through controlling ligand availability and fibrillar structure within another polymer that regulates bulk mechanical properties. These results provide new insight into the role of fibrin presentation and crosslinking density in alginate and address critical issues for hydrogel development. Moreover, these data emphasize the connection between gel stiffness and mesh size to influence cell response. This material may find broad utility for studying cell responses in hydrogels with different stiffnesses and as a cell transplantation vehicle.

## DATA AVAILABILITY STATEMENT

Data supporting the findings of this manuscript are available from the corresponding author upon reasonable request.

## ACKNOWLEDGEMENTS

Research reported in this publication was supported by National Institute of Dental and Craniofacial Research of the National Institutes of Health under award number R01 DE025475 and R01 DE025899 (JKL). CEV was supported by the NHLBI Training Program in Basic and Translational Cardiovascular Science (T32 HL086350). TGF was supported by the American Heart Association Postdoctoral Fellowship (19POST34460034). The content is solely the responsibility of the authors and does not necessarily represent the official views of the National Institutes of Health. We are grateful to Prof. Eduardo Silva for cord blood-derived endothelial cells.

